# Development of a Novel Japanese Eel Myoblast Cell Line for Application in Cultured Meat Production

**DOI:** 10.1101/2024.08.10.607424

**Authors:** Daisuke Ikeda, Yui Otsuka, Nobuhiro Kan-no

**Author notes:** Corresponding author. E-mail address (D. Ikeda).

## Abstract

The present study investigates the isolation, analysis, and characterization of primary cultured cells derived from the muscle tissue of Japanese eel (*Anguilla japonica*), culminating in establishing a spontaneously immortalized myoblast cell line, JEM1129. We isolated satellite cells from eel muscle tissue to establish a foundation for cultured eel meat production. While initial cell cultures contained myoblasts, continued passaging led to a decline in myoblast characteristics and an increase in fibroblast-like cells. RNA-Seq and RT-qPCR analyses showed significant downregulation of well-established markers for satellite cells and myoblasts, such as pax7a and myoD, over successive passages, highlighting a loss of myoblastic traits. Single-cell cloning was employed to overcome this challenge and maintain myoblast purity, leading to the successful creation of the JEM1129 cell line. These JEM1129 cells demonstrated enhanced expression of myoblast marker genes, exceeding the initial primary culture cell population. The cells showed strong myotube formation, particularly when cultured in a differentiation medium, indicating their robust potential for muscle development. The JEM1129 cell line represents a significant advancement in the cultivation of eel muscle cells, offering a promising avenue for cultured meat production. The findings contribute to a deeper understanding of muscle cell biology and provide valuable insights into using fish-derived myoblasts for cultured meat production.

**Highlights:** Single-cell cloning established a stable Japanese eel myoblast line, JEM1129.

Single-cell cloning aids in establishing spontaneously immortalized fish myoblasts.

JEM1129 demonstrates robust myogenic differentiation, crucial for cultured meat.

Potential for sustainable eel meat production reducing wild eel depletion.

## 1. Introduction

Cultured meat, also known as lab-grown or in vitro meat, has emerged as a promising alternative to traditional livestock farming, offering potential solutions to environmental, ethical, and food security concerns. This innovative approach involves cultivating animal cells in a controlled laboratory environment to produce meat products without animal slaughter [1–3]. While significant progress has been made in developing cultured meat from terrestrial animals, there is growing interest in exploring aquatic species as potential sources for cultured seafood [4–6].

The Japanese eel (*Anguilla japonica*) is a high-end food fish prized in Japan for its delicious taste. Most of the eels in the market are farmed, and the farming process begins with the collection of natural glass eels, which are juvenile eels in their early developmental stage [7]. However, wild eel populations, designated as endangered species due to their significant depletion [8], urgently require research aimed at their sustainable use. Cultured eel meat could provide a sustainable alternative to farmed eels, which rely on wild-caught glass eels, helping to alleviate pressure on wild populations while meeting consumer demand. This study aims to establish a foundation for producing cultured Japanese eel meat by isolating and characterizing muscle-derived cells, focusing on satellite cells/myoblasts. Satellite cells (myosatellite cells) are stem cells responsible for forming skeletal muscle and play a crucial role in muscle growth and regeneration. Upon activation, satellite cells become muscle progenitor cells, known as myoblasts, and proliferate through multiple cell divisions [9,10]. We isolated satellite cells from Japanese eel muscle tissue and attempted to generate myoblast strains from these cells. As a result, we could regularly isolate myoblast cells; however, a notable issue arose—the proportion of myoblast cells decreased over time and transitioned to a fibroblast-like population with continued passaging. To elucidate the molecular mechanism of this change, RNA-Seq analysis was conducted to investigate the variation in gene expression within the cell population during passaging, and qPCR primers were designed based on these findings to evaluate myoblasts. Furthermore, capitalizing on the characteristic tendency of fish cells to undergo spontaneous immortalization, we attempted single-cell cloning in the early stages of culture to establish cell lines derived from a single myoblast.

## 2. Materials and methods

### 2.1. Japanese eel care

Glass eel specimens of the Japanese eel *Anguilla japonica* were obtained from commercial suppliers, maintained at 28 °C, and were hand-fed daily with regular commercial feed. All animal experiments complied with the regulations set by the Animal Experimentation Committee of the School of Marine Biosciences, Kitasato University.

### 2.2. Isolation and cultivation of myoblasts

Skeletal muscle tissue was isolated from Japanese eels raised from glass eels and grown to 0.6-1.7 g as juveniles. The isolated muscle tissue was cut into small pieces using scissors and then washed three times in L-15 medium (Thermo Fisher Scientific, MA, U.S.A.) containing 5% Antibiotic-Antimycotic Mixed Stock Solution (Nacalai Tesque, Kyoto, Japan). The tissue pieces were then placed in 50 mL tubes with 3.3 mg/mL of Collagenase Type IV (Worthington, NJ, U.S.A) and digested for 1 hour at room temperature while rotating in a roller mixer. After collagenase digestion, Trypsin (MP biomedicals, CA, U.S.A.) was added at a final concentration of 1 mg/mL and digested for another 20 minutes. The obtained cells were strained through 70 and 40 μm cell strainers and suspended in growth medium (GM), L-15 medium containing 20% fetal bovine serum (FBS) (Sigma Aldrich, MA, U.S.A.), 1% Antibiotic-Antimycotic Mixed Stock Solution, 1% MEM Non-essential Amino Acids Solution (FUJIFILM Wako Pure Corporation Chemical, Osaka, Japan), and 20 ng/mL of bFGF (KAC, Kyoto, Japan). The suspended cells were then seeded into 6-well plates pre-coated with Easy iMatrix-511 (MATRIXOME, Osaka, Japan) and cultured at 28 °C.

### 2.3. RNA-Sequencing (RNA-Seq)

To comprehensively understand the changes in gene expression, we performed RNA-Seq on the cell population isolated from muscle tissue. We prepared RNA-Seq libraries from cells with varying passage numbers using the ISOSPIN Cell & Tissue RNA (Nippon Gene, Tokyo, Japan), the KAPA mRNA Capture Kit (Roche, Basel, Switzerland), and the MGIEasy RNA Directional Library Prep Set (MGI Tech Co., Ltd., Shenzhen, China). We sequenced these libraries with 150 bp paired-end reads using the DNBSEQ-G400RS platform. Genome-Lead Corp. (Kagawa, Japan) conducted the RNA-Seq, and we submitted the processed reads to the DDBJ Sequence Read Archive (DRA) under accession number DRA017096.

### 2.4. Mapping of sequence reads, differential expression analysis, and gene enrichment analysis

Using the Japanese eel genome sequence (ASM2516954v1) [11] as a reference, read sequences obtained from primary myoblast populations (n=2) derived from Japanese eel muscle tissue at 1, 8, and 16 passages were mapped using HISAT2 [12]. A separately published gene annotation file (Anguilla_japonica.gene.0224.gff) [11] and Cuffdiff [13] were used to identify genes whose expression levels significantly decreased with an increasing number of passages (Supplementary Data 1). To focus on genes whose expression decreased with the number of passages in this study, a group of genes whose expression decreased more than fourfold compared to the initial culture period was extracted. For the 29,982 genes predicted by the gene annotation file, we conducted an independent BLAST search against the Ensembl zebrafish database and identified 26,772 homologous genes. Biological interpretation of the list of genes with altered expression levels was performed by enrichment analysis using Metascape [14].

### 2.5. RT-qPCR analysis

To evaluate muscle cell characteristics using RT-qPCR, we selected muscle- and adipocyte-related genes that exhibited passage-dependent downregulation in the RNA-Seq analysis and designed qPCR primers targeting these genes using Primer-BLAST. Cells were cultured in 12-well plates until reaching 80-90% confluence. We then extracted total RNA using ISOGEN II (Nippon Gene) and synthesized single-stranded cDNA using ReverTra Ace® qPCR RT Master Mix (TOYOBO, Osaka, Japan). We selected GAPDH as the reference gene for relative quantification [15]. Table S1 lists the primers used in this study. We performed qPCR reactions using THUNDERBIRD® Next SYBR™ qPCR Mix (Toyobo) and quantified the gene expression levels using the ⊿⊿Ct method.

The statistical analysis was performed with R Statistical Software (version 4.2.2). We analyzed relative expression levels using a one-way ANOVA, followed by a Tukey-Kramer post-hoc test. A p-value below 0.05 indicated a statistically significant difference.

### 2.6. Single-cell cloning

We counted the primary myoblast populations derived from Japanese eel muscle at passages 1-3. We adjusted them to a concentration of approximately 0.5 cells per 100 µL, ensuring less than one cell per well. We seeded this cell suspension into 96-well plates at 100 µL per well. Over time, we observed the plates and trypsinized wells showing sufficient cell growth to detach the cells, which we then passed into 24-well plates. After allowing for adequate proliferation, we examined the cell morphology and selected clones exhibiting a myoblast-like appearance—characterized by an elongated spindle shape—and transferred them to 6-well plates. We further expanded the cultures by sequentially transitioning to T25 and T75 flasks. We performed time-lapse imaging using a Lux2 microscope (Axion BioSystems, GA, U.S.A.). We preserved the established cell lines in liquid nitrogen using Cell Reservoir One (with DMSO) cryopreservation solution (Nacalai Tesque).

### 2.7. Immunocytochemistry

JEM1129 cells were cultured in a Slide & Chamber 4 Wells (Watson, Tokyo, Japan) in GM until they reached 90% confluence. The medium was then changed to a differentiation medium (DM) consisting of L-15 medium containing 2% horse serum (HS) (Sigma Aldrich) and 1% Antibiotic-Antimycotic Mixed Stock Solution to promote differentiation into myotube cells. The myotubes were fixed with 4% paraformaldehyde for 5 minutes, permeabilized in PBS containing 0.1% Triton X-100 for 5 minutes, and blocked with 3% bovine serum albumin (BSA) for 10 minutes. The primary antibody used was the anti-myosin heavy chain antibody MF20 (Developmental Studies Hybridoma Bank, IA, U.S.A.) at a 1:50 dilution. The immunoreaction with the primary antibody was performed overnight at 4°C, followed by incubation with anti-mouse IgG DyLight 549 (ROCKLAND, PA, U.S.A.) at a 1:1000 dilution for 1 hour at room temperature. Nuclei were stained with Hoechst 33342 (Sigma Aldrich). Stained cells were visualized using a DMI 4000B fluorescence microscope (Leica, Wetzlar, Germany).

## 3. Results and Discussion

### 3.1. Isolation and Analysis of Primary Cultured Cells from Muscle Tissue

Approximately 200 mg of muscle tissue was enzyme digested and seeded in equal amounts into laminin-coated 6-well plates. After one week of culture, the cells proliferated to around 5.0 × 10^5^ cells per well, so the culture was expanded into T25 flasks, and successive passages were continued. Immediately after expansion from the 6-well plates into T25 flasks (first passage), the cell population contained many spindle-shaped small cells that appeared to be myoblasts (Fig. 1). After the primary culture cells reached subconfluence during the first to third passages, the medium was switched from GM to DM and the cells were cultured for one week, during which many elongated myotube-like cells were observed (Fig. S1 A, B). When the differentiated cells were detached from the culture vessel using trypsin, numerous elongated cells resembling mature myocytes were observed (Fig. S1 C, D).

**Figure 1.**
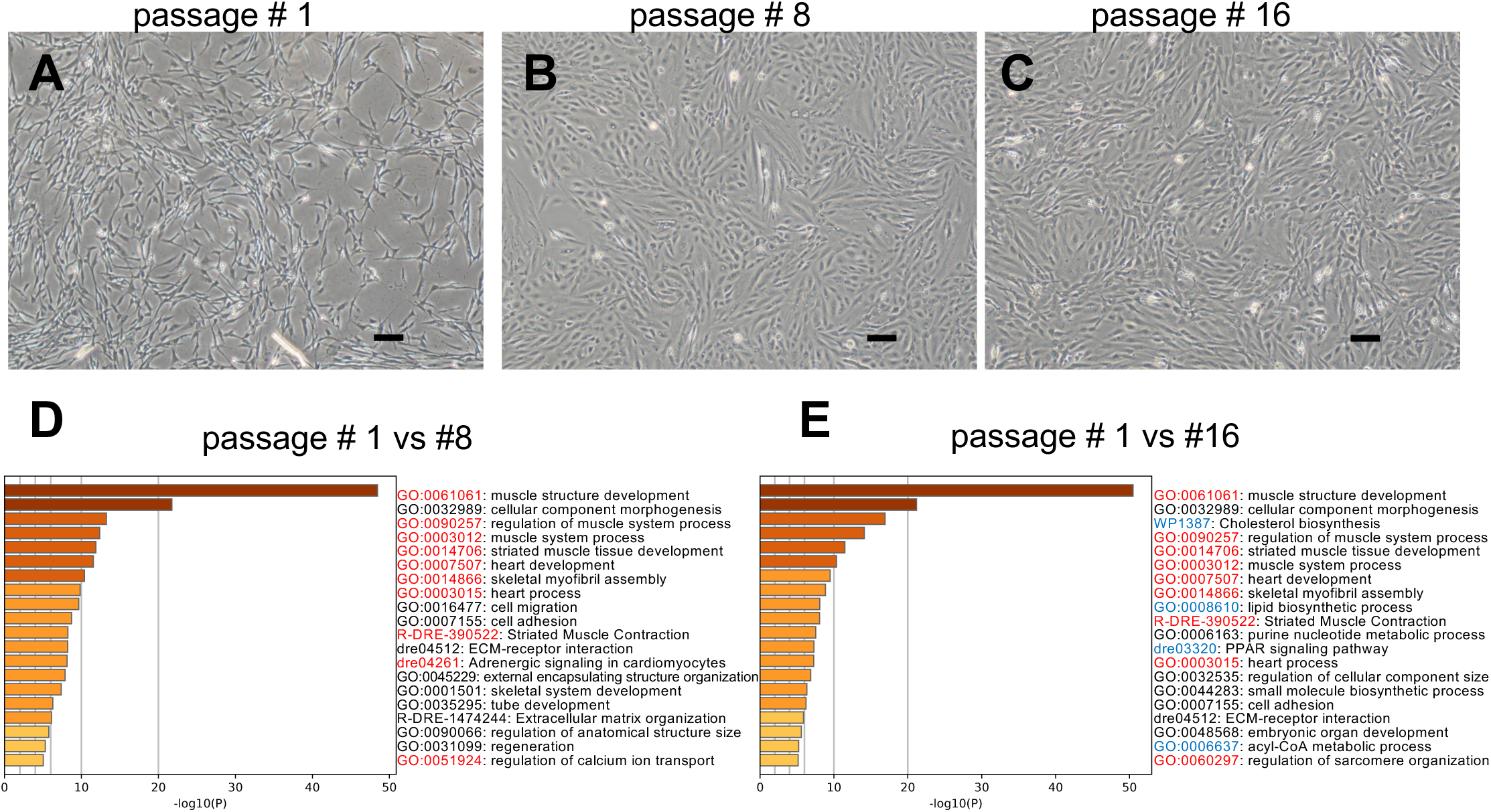
Changes in primary cultured cells derived from Japanese eel muscle tissue with increasing passages. (A-C) Phase contrast view (scale bars: 100 µm). With continued passaging, the percentage of myoblast-like cells (A) decreased, while the percentage of fibroblast-like cells increased (B and C). (D, E) Bar graph of enriched terms across input gene lists. Genes with a 4-fold or greater decrease in expression in the cell population at the beginning of culture and at the 8^th^ (D) and 16th (E) passages were extracted and analyzed. The IDs considered characteristic of myocytes and adipocytes are shown in red and blue, respectively.

However, with continued passaging, the percentage of myoblast-like cells decreased while the percentage of fibroblast-like cells increased (Fig. 1B, C). Inducing differentiation in these cells did not result in myotube formation. When isolating myoblasts from muscle tissue, it is crucial to minimize fibroblast contamination [16]. This is because fibroblasts are fast-dividing and easy to culture [17], and even a tiny percentage of fibroblasts can outgrow and replace a population initially abundant in myoblasts. Satellite cells isolated from muscle tissue for cultured meat production can be purified through cell sorting using specific molecular markers in pigs [18,19] and cattle [reviewed in [1]] to prevent fibroblast contamination before culture. However, in Japanese eels, purification by cell sorting is not feasible due to the lack of established molecular markers for satellite cell isolation.

Previous studies [18,19] have reported that even satellite cells purified to high purity by cell sorting lose their stem cell characteristics due to decreased expression of marker genes such as Pax7 with passaging. Consistent with this, we observed that cell populations isolated from Japanese eel muscle tissue lost their myoblastic characteristics with passaging, as evidenced by changes in cell morphology (Fig. 1A-C). To elucidate the molecular mechanism of this change, we analyzed the expression levels of genes that characterize myoblasts by RNA-Seq analysis. We found that the expression levels of 1320 and 1382 genes decreased more than 4-fold after 8 and 16 passages, respectively, with 1270 and 1332 of these genes having zebrafish orthologs, respectively (Supplementary Data 2). An enrichment analysis using Metascape was performed on these passage-dependently down-regulated genes to elucidate their functional implications. The enrichment analysis revealed distinct patterns of change during cell passaging. At 8 passages, a notable decrease in muscle-related gene expression was observed (Fig. 1D, Supplementary Data 3). By 16 passages, the reduction extended to both muscle-related and lipid-related genes (Fig. 1E, Supplementary Data 4). These findings suggest that the initially isolated cell population comprised both myocytes and adipocytes.

To characterize adipocytes, we selected six genes classified under GO:0008610 (lipid biosynthetic process) as marker genes and designed RT-qPCR primers for these targets. We designed RT-qPCR primers for myoblast characterization for five genes categorized under GO:0061061 (muscle structure development). We also included primers for Pax7 and Cdh15 (M-cadherin), identified as differentially expressed genes in our RNA-Seq analysis, and well-established markers for satellite cells and myoblasts [9,10,20]. RT-qPCR analysis confirmed the RNA-Seq data, showing that the expression levels of target genes decreased as the number of passages increased (Figs. 2 and 3). Similar to the RNA-Seq results, the expression levels of target genes for myocytes decreased significantly at 8 passages, with no significant difference between cells after 8 and 16 passages (Fig. 2). For three of the six adipocyte target genes, fasn, ebp, and fabp1b.1, expression levels in cells after 8 passages were significantly higher than those after 16 passages, supporting the RNA-Seq results (Fig. 3). This strongly suggests that the RT-qPCR method using these primers is effective for the evaluation of adipocytes as well as myocytes in Japanese eel. While muscle cells have been the primary focus of cultured meat research, adipocytes are also critical components of muscle tissue. The adipocytes are particularly important for eels, commonly served as “glaze-grilled eel,” where the fat content in the meat significantly affects the taste. The adipocyte primers designed in this study will be highly valuable for future research on the isolation and study of adipocytes from Japanese eel.

**Figure 2.**
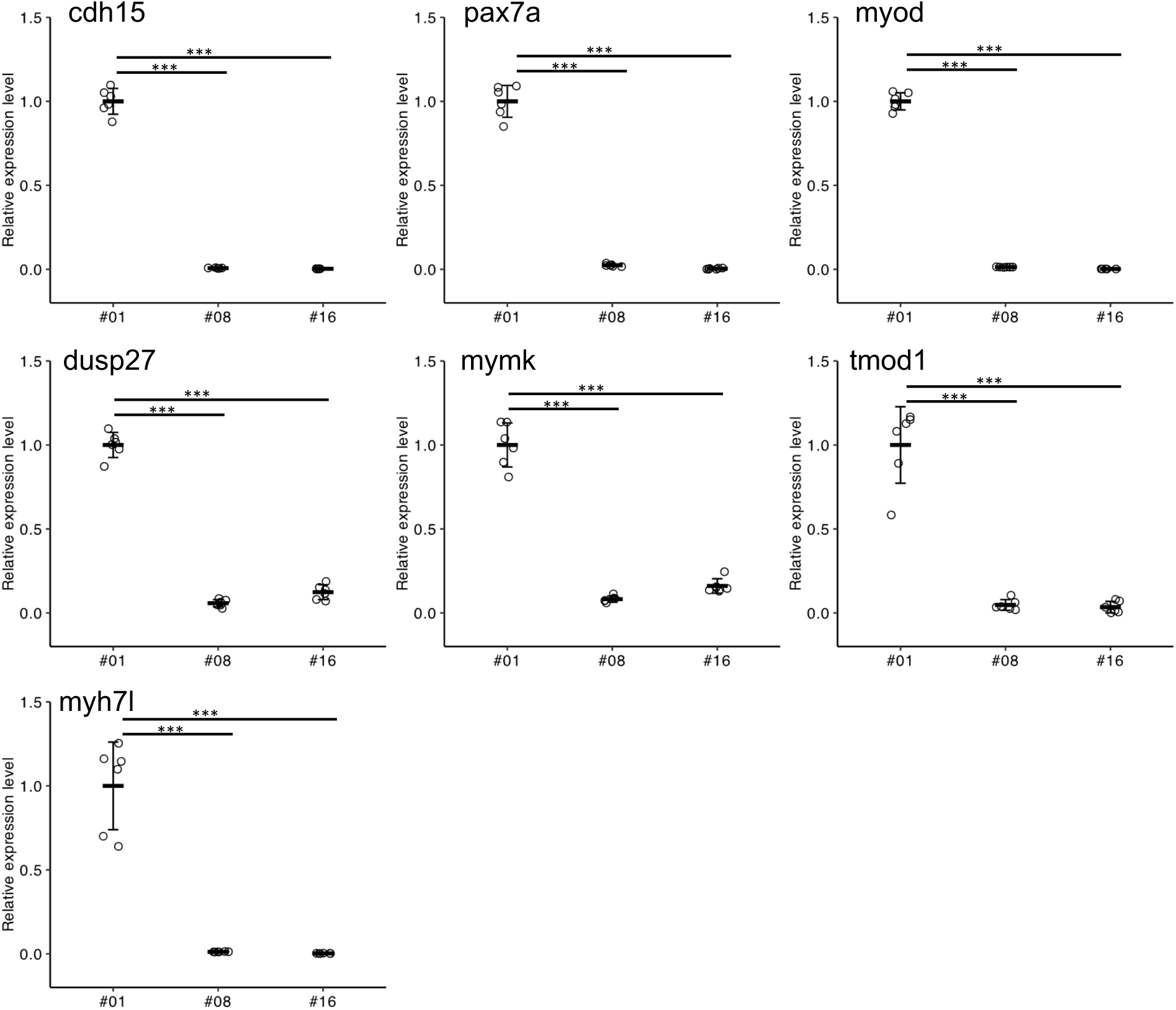
The relationship between passage number and the expression levels of myocyte marker genes in primary cultured myoblasts populations. The relative expression levels were normalized using GAPDH as a reference and presented as a ratio to the primary culture cells of the first passage. The expression levels are presented as means ± standard deviations (n=6). In the post-hoc test, asterisks indicate statistical significance in comparisons between expression levels under different conditions (***: p < 0.001).

**Figure 3.**
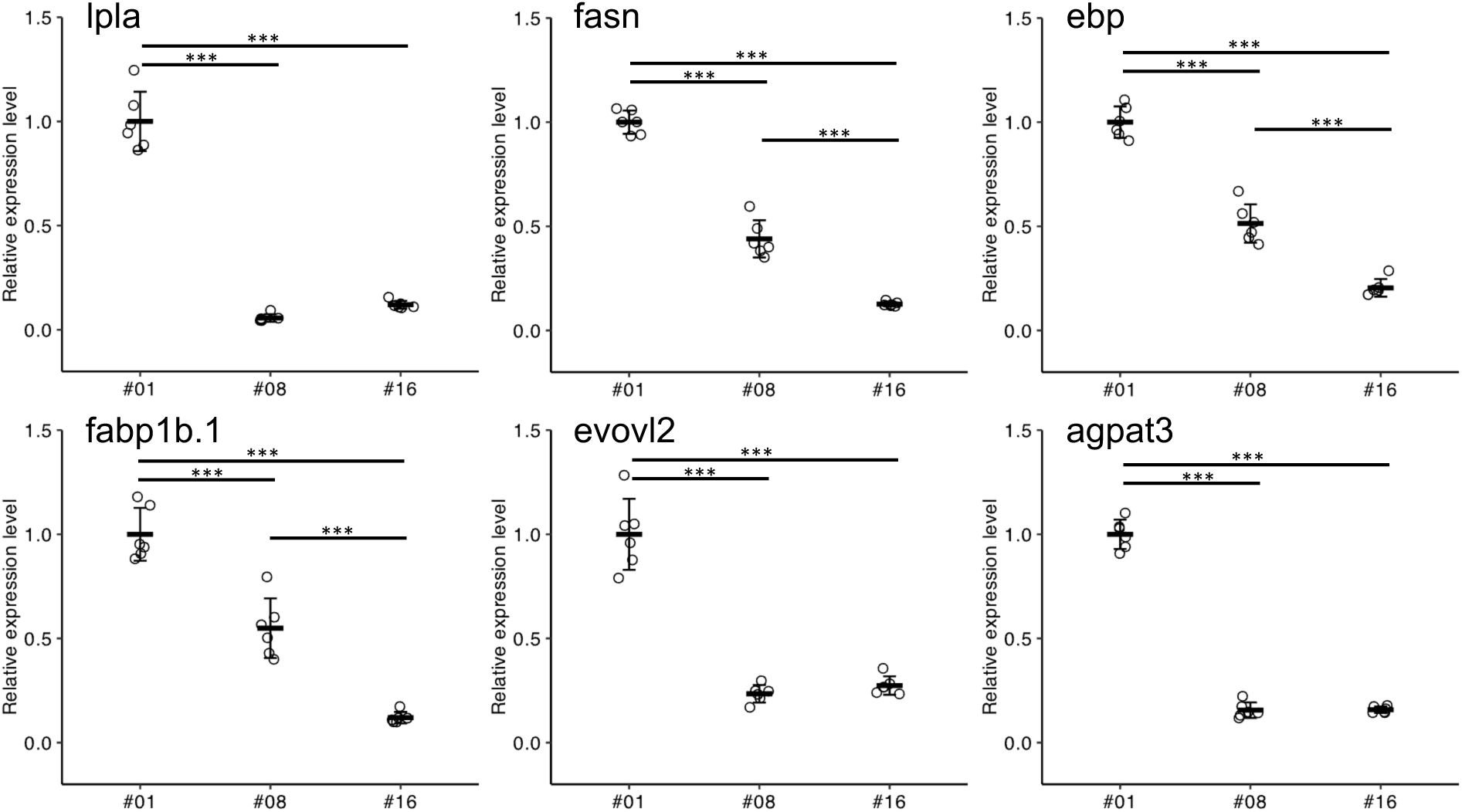
The relationship between passage number and the expression levels of adipocyte marker genes in primary cultured myoblasts populations. The relative expression levels were normalized using GAPDH as a reference and presented as a ratio to the primary culture cells of the first passage. The expression levels are presented as means ± standard deviations (n=6). In the post-hoc test, asterisks indicate statistical significance in comparisons between expression levels under different conditions (***: p < 0.001).

### 3.2. Establishment of Japanese Eel Myoblast Cell Line

Myoblast cell lines have been established in fish from species such as black rockfish [21], Atlantic mackerel [22], and olive flounder [23]. These reports indicated that myoblast cell lines were generated by culturing cells isolated from muscle tissue. However, they did not specify whether satellite cells/myoblasts were actively purified during the process. Initially, we also conducted several experiments expecting to obtain similar cell types by continuously passaging cells isolated from the muscle tissue of Japanese eel. However, as described above, the myoblast characteristics inevitably diminished and were progressively replaced by a fibroblast-like population as passaging continued.

Unlike most mammals, fish grow continuously without showing signs of aging [24,25]. This ongoing growth requires their cells to keep dividing and their tissues to repair themselves even when fully developed. Notably, fish cell cultures tend to multiply indefinitely without turning cancerous, suggesting fish cells can achieve immortality without becoming malignant [26,27]. Although the exact cause of fish cells’ susceptibility to spontaneous immortality is unknown, Futami et al. suggested that the lack of the p16^INK4a^/ARF, a cell cycle arrest factor and marker gene for senescent cells, locus in the fish genome may be a factor in spontaneous immortality [27]. Therefore, given the characteristic tendency of fish cells to undergo spontaneous immortalization, we attempted single-cell cloning during the early stages of culture to establish cell lines derived from a single myoblast.

Single-ce4l cloning of the cell population immediately after isolation from Japanese eel muscle tissue yielded several types of cloned cells. Among these, clones exhibiting a myoblast-like morphology were initially observed as individual cells immediately after seeding (Fig. 4A). Spontaneously immortalized myoblasts were actively dividing (Video S1), and their doubling time was about 30-36 hours. As these cells proliferated and reached confluence—covering the adhesive surface of the culture vessel—they spontaneously fused to form elongated myotube cells (Fig. 4B). Upon detaching the cell population in this state using trypsin, numerous elongated fibrous myotube cells were observed alongside spherical undifferentiated myoblasts (Fig. 4C, arrowheads). We designated the myoblast cell line derived from a single cell as Japanese Eel Myoblast 1129 (JEM1129).

**Figure 4.**
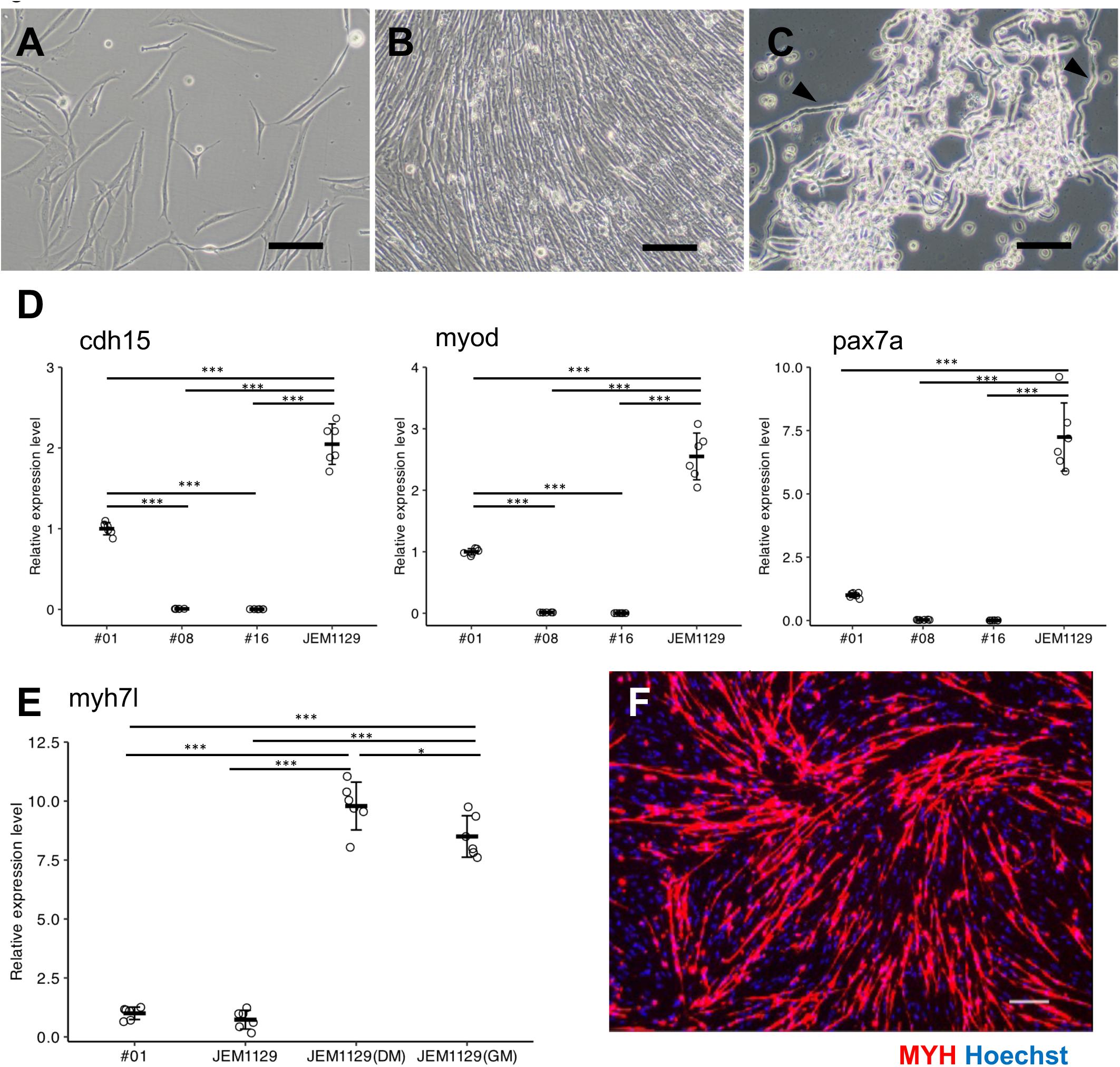
Established Japanese Eel Myoblast Cell Line JEM1129. (A-C) Phase contrast images (scale bars: 100 µm). (A) Cells before reaching confluence. (B) Cells after reaching confluence. (C) Upon detachment of the confluent cell population using trypsin, numerous fibrous myotubes were observed (indicated by arrowheads). (D, E) Comparison of myocyte marker gene expression levels in primary cultured myoblasts and JEM1129. The expression levels are presented as means ± standard deviations (n=6). In the post-hoc test, asterisks indicate statistical significance in comparisons between different conditions (*: p < 0.05, ***: p < 0.001). (D) All well-established markers for satellite cells and myoblasts exhibit higher expression levels in JEM1129 than in the first passage’s primary culture cells. (E) The expression levels of myosin heavy chain genes before muscle differentiation are similar in JEM1129 and the primary culture cells of the first passage (#1 and JEM1129). When the differentiation medium induces muscle differentiation, intracellular myosin heavy chain gene expression increases (JEM1129(DM)). This increase occurs even without replacement with a differentiation medium (JEM1129(GM)). (F) Immunofluorescence staining for myosin heavy chain (MYH) and cell nuclei (Hoechst). JEM1129 was induced into myotubes using a differentiation medium. Scale bar: 100 µm.

As shown in Figure 2, the expression levels of myoblast marker genes in the cell population derived from muscle tissue decrease significantly at passage 8. However, the cloned myoblast line JEM1129 displayed similar or higher expression levels of myocyte marker genes than the initial cell population (Figs. 4D and S2). Notably, the expression levels of pax7a, myoD, and cdh15, which are well-established markers for satellite cells and myoblasts [9,10,20], were more than twice as high as in the initial cell population, with pax7a showing a 7-fold increase (Fig. 4D).

After JEM1129 cells reached subconfluence, the medium was switched from GM to DM and cultured for one week. The expression level of myosin heavy chain, the established marker gene for muscle differentiation [9,10], was compared by RT-qPCR between JEM1129 cells before reaching subconfluence and primary culture cells, and the expression level was about 10-fold higher (Fig. 4E), strongly suggesting that the cells differentiated into muscle. Immunofluorescence staining results also confirmed the strong expression of myosin heavy chains in JEM1129 cells, in which DM-induced muscle differentiation (Fig. 4F). However, RT-qPCR results showed that myosin heavy chain expression was also increased in cells cultured in GM without switching medium to DM (Fig. 4E). However, the expression level was significantly lower than in cells switched to DM (Fig. 4E).

In conclusion, the present study presents the successful isolation and characterization of a novel spontaneously immortalized Japanese eel myoblast cell line, JEM1129. This cell line was achieved through single-cell cloning of a primary muscle tissue cell population, overcoming the challenge of fibroblast overgrowth and loss of myoblast characteristics observed during conventional passaging. By attempting single-cell cloning of fish myoblasts, characterized by their tendency to immortalize spontaneously in the early stages of culture, myoblast cell lines are expected to be established in many fish species. The novel myoblast cell line, JEM1129, presents significant potential for application in cultured Japanese eel meat production. Future studies should explore the scalability of culturing these cells and their differentiation into mature muscle tissue in bioreactors. Additionally, integrating adipocytes into these cultures, as they play a critical role in the taste of eel meat, could lead to more refined and commercially viable cultured meat products.

## Funding

This study is financially supported by JSPS KAKENHI Grant Number 20K06232 and the Ichimasa Kamaboko Co., Ltd. research grant.

## Author Contributions

Conceived and designed the experiments: DI, YO, and NK. Performed the experiments: DI and YO Analyzed the data: DI and YO. Wrote the manuscript: DI.

## Availability of data and materials

The data and materials are available according to request.

## Declaration of Competing Interest

The authors declare no conflict of interest.

## Acknowledgments

We especially mention Dr. Hiroaki Chiba for his contribution to glass eel breeding. We are very grateful to Dr. Yoji Nakamura for his help in the RNA-Seq data analysis.

## Notes

### Competing Interest Statement

The authors have declared no competing interest.

### Summary of Updates

Change the author name Nobushiro =>; Nobuhiro

## References

[1] M. Olenic, C. Deelkens, E. Heyman, E. De Vlieghere, X. Zheng, J. van Hengel, C. De Schauwer, B. Devriendt, S. De Smet, L. Thorrez, Review: Livestock cell types with myogenic differentiation potential: Considerations for the development of cultured meat, Animal (2024) 101242. 10.1016/j.animal.2024.101242.

[2] N. Treich, Cultured Meat: Promises and Challenges, Environ Resource Econ 79 (2021) 33–61. 10.1007/s10640-021-00551-3.

[3] M.J. Post, Cultured beef: medical technology to produce food, Journal of the Science of Food and Agriculture 94 (2014) 1039–1041. 10.1002/jsfa.6474.

[4] R. Ovissipour, X. Yang, Y.T. Saldana, D.L. Kaplan, N. Nitin, A. Shirazi, B. Chirdon, W. White, B. Rasco, Cell-based fish production case study for developing a food safety plan, Heliyon 10 (2024) e33509. 10.1016/j.heliyon.2024.e33509.

[5] M. Goswami, Y. Belathur Shambhugowda, A. Sathiyanarayanan, N. Pinto, A. Duscher, R. Ovissipour, W.S. Lakra, R. Chandragiri Nagarajarao, Cellular Aquaculture: Prospects and Challenges, Micromachines 13 (2022) 828. 10.3390/mi13060828.

[6] M. Goswami, B.S. Yashwanth, V. Trudeau, W.S. Lakra, Role and relevance of fish cell lines in advanced in vitro research, Mol Biol Rep 49 (2022) 2393–2411. 10.1007/s11033-021-06997-4.

[7] L.T.N. Heinsbroek, A review of eel culture in Japan and Europe, Aquaculture Research 22 (1991) 57–72. 10.1111/j.1365-2109.1991.tb00495.x.

[8] D.M.P. Jacoby, J.M. Casselman, V. Crook, M.-B. DeLucia, H. Ahn, K. Kaifu, T. Kurwie, P. Sasal, A.M.C. Silfvergrip, K.G. Smith, K. Uchida, A.M. Walker, M.J. Gollock, Synergistic patterns of threat and the challenges facing global anguillid eel conservation, Global Ecology and Conservation 4 (2015) 321–333. 10.1016/j.gecco.2015.07.009.

[9] E.N. Engquist, P.S. Zammit, The Satellite Cell at 60: The Foundation Years, Journal of Neuromuscular Diseases 8 (2021) S183–S203. 10.3233/JND-210705.

[10] M. Schmidt, S.C. Schüler, S.S. Hüttner, B. von Eyss, J. von Maltzahn, Adult stem cells at work: regenerating skeletal muscle, Cell. Mol. Life Sci. 76 (2019) 2559–2570. 10.1007/s00018-019-03093-6.

[11] H. Wang, H.T. Wan, B. Wu, J. Jian, A.H.M. Ng, C.Y.-L. Chung, E.Y.-C. Chow, J. Zhang, A.O.L. Wong, K.P. Lai, T.F. Chan, E.L. Zhang, C.K.-C. Wong, A Chromosome-level assembly of the Japanese eel genome, insights into gene duplication and chromosomal reorganization, GigaScience 11 (2022) giac120. 10.1093/gigascience/giac120.

[12] D. Kim, J.M. Paggi, C. Park, C. Bennett, S.L. Salzberg, Graph-based genome alignment and genotyping with HISAT2 and HISAT-genotype, Nat Biotechnol 37 (2019) 907–915. 10.1038/s41587-019-0201-4.

[13] C. Trapnell, A. Roberts, L. Goff, G. Pertea, D. Kim, D.R. Kelley, H. Pimentel, S.L. Salzberg, J.L. Rinn, L. Pachter, Differential gene and transcript expression analysis of RNA-seq experiments with TopHat and Cufflinks, Nat Protoc 7 (2012) 562–578. 10.1038/nprot.2012.016.

[14] Y. Zhou, B. Zhou, L. Pache, M. Chang, A.H. Khodabakhshi, O. Tanaseichuk, C. Benner, S.K. Chanda, Metascape provides a biologist-oriented resource for the analysis of systems-level datasets, Nat Commun 10 (2019) 1523. 10.1038/s41467-019-09234-6.

[15] J. Gu, S. Dai, H. Liu, Q. Cao, S. Yin, K.P. Lai, W.K.F. Tse, C.K.C. Wong, H. Shi, Identification of immune-related genes in gill cells of Japanese eels (*Anguilla japonica*) in adaptation to water salinity changes, Fish & Shellfish Immunology 73 (2018) 288–296. 10.1016/j.fsi.2017.12.026.

[16] D.C. Stephens, M. Mungai, A. Crabtree, H.K. Beasley, E. Garza-Lopez, L. Vang, K. Neikirk, Z. Vue, N. Vue, A.G. Marshall, K. Turner, J. Shao, B. Sarker, S. Murray, J.A. Gaddy, J. Davis, S.M. Damo, A.O. Hinton, Protocol for isolating mice skeletal muscle myoblasts and myotubes via differential antibody validation, STAR Protocols 4 (2023) 102591. 10.1016/j.xpro.2023.102591.

[17] I.R. Fernandes, F.B. Russo, G.C. Pignatari, M.M. Evangelinellis, S. Tavolari, A.R. Muotri, P.C.B. Beltrão-Braga, Fibroblast sources: Where can we get them?, Cytotechnology 68 (2016) 223–228. 10.1007/s10616-014-9771-7.

[18] H. Zhu, Z. Wu, X. Ding, M.J. Post, R. Guo, J. Wang, J. Wu, W. Tang, S. Ding, G. Zhou, Production of cultured meat from pig muscle stem cells, Biomaterials 287 (2022) 121650. 10.1016/j.biomaterials.2022.121650.

[19] S. Ding, F. Wang, Y. Liu, S. Li, G. Zhou, P. Hu, Characterization and isolation of highly purified porcine satellite cells, Cell Death Discov. 3 (2017) 1–11. 10.1038/cddiscovery.2017.3.

[20] A.J. Goel, M.-K. Rieder, H.-H. Arnold, G.L. Radice, R.S. Krauss, Niche Cadherins Control the Quiescence-to-Activation Transition in Muscle Stem Cells, Cell Reports 21 (2017) 2236–2250. 10.1016/j.celrep.2017.10.102.

[21] X. Kong, X. Wang, M. Li, W. Song, K. Huang, F. Zhang, Q. Zhang, J. Qi, Y. He, Establishment of myoblast cell line and identification of key genes regulating myoblast differentiation in a marine teleost, Sebastes schlegelii, Gene 802 (2021) 145869. 10.1016/j.gene.2021.145869.

[22] M.K. Saad, J.S.K. Yuen, C.M. Joyce, X. Li, T. Lim, T.L. Wolfson, J. Wu, J. Laird, S. Vissapragada, O.P. Calkins, A. Ali, D.L. Kaplan, Continuous fish muscle cell line with capacity for myogenic and adipogenic-like phenotypes, Sci Rep 13 (2023) 5098. 10.1038/s41598-023-31822-2.

[23] S. Krishnan, S. Ulagesan, J. Cadangin, J.-H. Lee, T.-J. Nam, Y.-H. Choi, Establishment and Characterization of Continuous Satellite Muscle Cells from Olive Flounder (Paralichthys olivaceus): Isolation, Culture Conditions, and Myogenic Protein Expression, Cells 12 (2023) 2325. 10.3390/cells12182325.

[24] C.E. Finch, Update on Slow Aging and Negligible Senescence – A Mini-Review, Gerontology 55 (2009) 307–313. 10.1159/000215589.

[25] D. Reznick, C. Ghalambor, L. Nunney, The evolution of senescence in fish, Mechanisms of Ageing and Development 123 (2002) 773–789. 10.1016/S0047-6374(01)00423-7.

[26] N.C. Bols, L.E.J. Lee, G.C. Dowd, Distinguishing between ante factum and post factum properties of animal cell lines and demonstrating their use in grouping ray-finned fish cell lines into invitromes, In Vitro Cell.Dev.Biol.-Animal 59 (2023) 41–62. 10.1007/s11626-022-00744-0.

[27] K. Futami, S. Sato, M. Maita, T. Katagiri, Lack of a *p16*INK4a*/ARF* locus in fish genome may underlie senescence resistance in the fish cell line, EPC, Developmental & Comparative Immunology 133 (2022) 104420. 10.1016/j.dci.2022.104420.

