## Supplementary material for "Development of a Novel Japanese Eel Myoblast Cell Line for Application in Cultured Meat Production": Fig. S1

### Slide 1
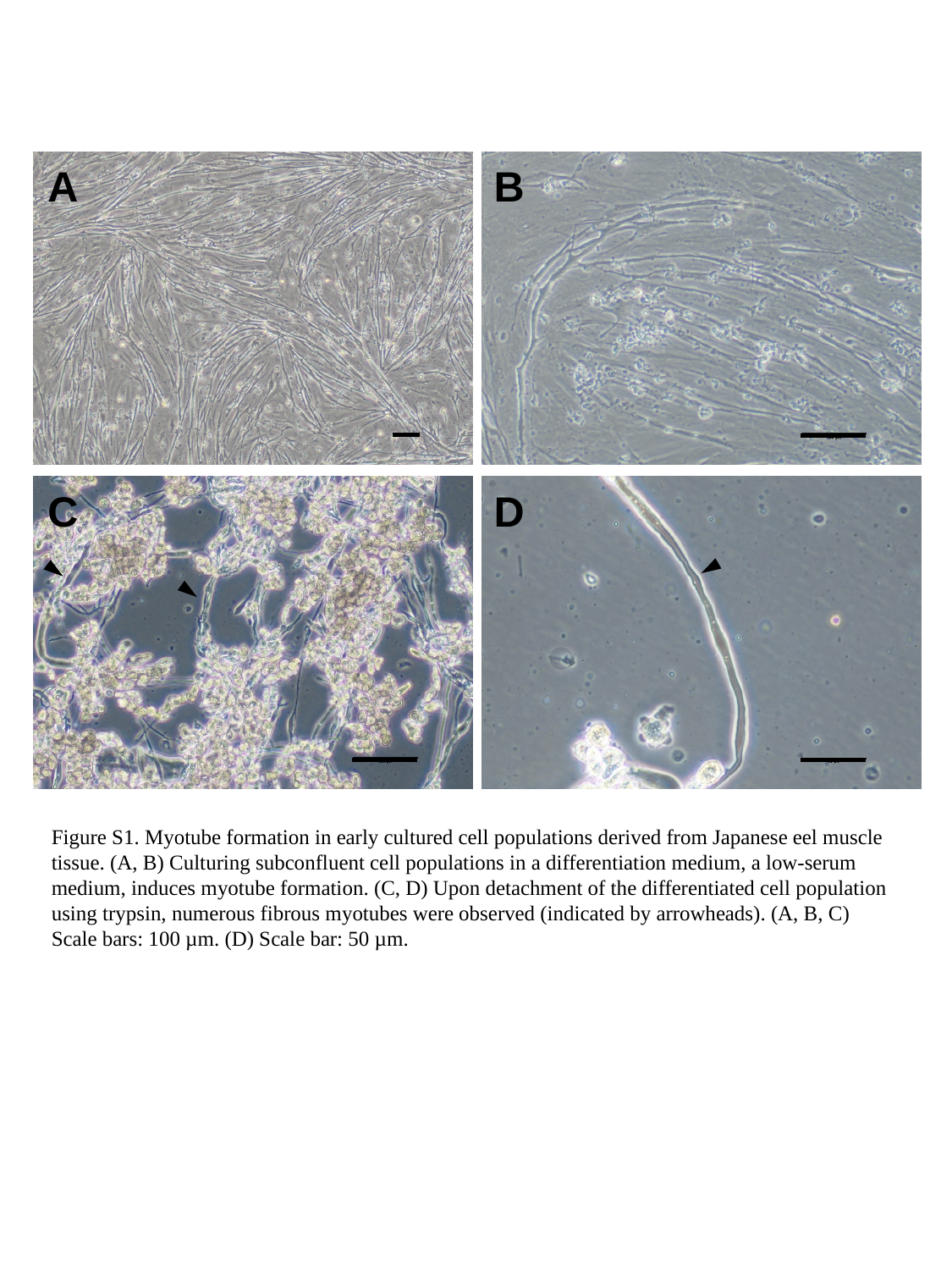

A
B
C
D
Figure S1. Myotube formation in early cultured cell populations derived from Japanese eel muscle tissue. (A, B) Culturing subconfluent cell populations in a differentiation medium, a low-serum medium, induces myotube formation. (C, D) Upon detachment of the differentiated cell population using trypsin, numerous fibrous myotubes were observed (indicated by arrowheads). (A, B, C) Scale bars: 100 µm. (D) Scale bar: 50 µm.
