## Supplementary material for "Development of a Novel Japanese Eel Myoblast Cell Line for Application in Cultured Meat Production": Fig. S2

### Slide 1
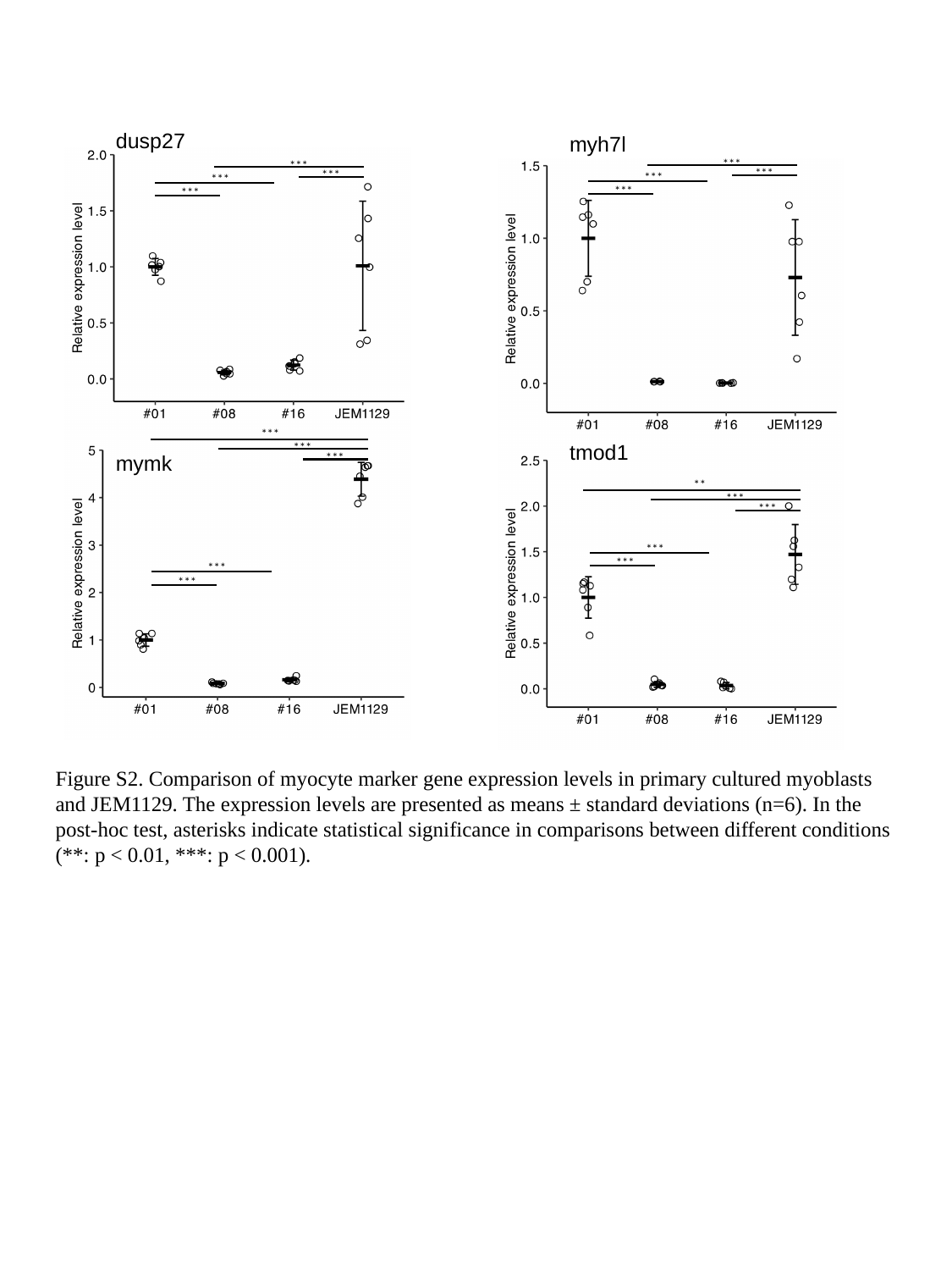

dusp27
myh7l
***
***
***
***
***
***
***
***
***
tmod1
***
***
mymk
**
***
***
***
***
***
***
Figure S2. Comparison of myocyte marker gene expression levels in primary cultured myoblasts and JEM1129. The expression levels are presented as means ± standard deviations (n=6). In the post-hoc test, asterisks indicate statistical significance in comparisons between different conditions (**: p < 0.01, ***: p < 0.001).
